# The neural determinants of abstract beauty

**DOI:** 10.1101/2022.07.13.499869

**Authors:** Samuel E. Rasche, Ahmad Beyh, Marco Paolini, Semir Zeki

**Author notes:** Contributed equally to this work (Samuel E. Rasche, Ahmad Beyh).

## Abstract

We have enquired into the neural activity which correlates with the experience of beauty aroused by abstract paintings consisting of arbitrary assemblies of lines and colours. During the brain imaging experiments, subjects rated abstract paintings according to aesthetic appeal. There was low agreement on the aesthetic classification of these paintings among participants. Univariate analyses revealed higher activity with higher declared aesthetic appeal in both the visual areas and the medial frontal cortex. Additionally, representational similarity analysis (RSA) revealed that the experience of beauty correlated with decodable patterns of activity in visual areas. These results are broadly similar to those obtained in previous studies on facial beauty. With abstract art, it was the involvement of visual areas implicated in the processing of lines and colours while with faces it was of visual areas implicated in the processing of faces. Both categories of aesthetic experience correlated with increased activity in medial frontal cortex. We conclude that the sensory areas participate in the selection of stimuli according to aesthetic appeal and that it is the co-operative activity between the sensory areas and the medial frontal cortex that is the basis for the experience of abstract visual beauty. Further, this co-operation is enabled by “experience dependent” functional connections, in the sense that currently the existence and high specificity of these connections can only be demonstrated during certain experiences.

## INTRODUCTION

What are the qualities in an object that arouse the sense of beauty, or what Clive Bell (1914) termed the “aesthetic emotion”? Yang et al. (2022) addressed this question by enquiring into the determinants of one of the most common sources of beauty, namely facial beauty. Features such as symmetry, proportion, and mathematically defined precise relationships between its constituent parts, have been posited by many, including leading artists such as Polykleitos and Leonardo Da Vinci, to be fundamental determinants of facial beauty, in the sense that without these biologically determined and inherited characteristics a face cannot be qualified as beautiful (Zeki, 2009; Zeki and Chén, 2020). But even though essential, these characteristics are not in themselves necessarily sufficient to render a face beautiful: there is, in addition, another, or other, unknown and mysterious characteristics that do so. What can they be? The Yang et al. (2022) study revealed that, when a face is perceived as beautiful and only then, a detectable neurobiological imprint in the form of particular decodable activity patterns emerges in sensory face-processing areas of the visual brain. The emergence of such decodable patterns correlates with the emergence of parallel decodable patterns in another part of the brain – the medial frontal cortex – in which activity always correlates with the experience of beauty, regardless of source (Ishizu and Zeki, 2011; Kawabata and Zeki, 2004). For faces at least, it is seemingly the joint activity of both components – the sensory on the one hand and the emotional on the other – that lies at the basis of the experience of beauty.

Facial beauty has been classified as belonging to the biological category of beauty (Zeki and Chén, 2020). In the present study, we ask the same question of artifactual beauty, that is, beauty generated by human agency. Many artists, including those belonging to the schools of Abstract Expressionism, Neo-Plasticism and Russian Constructivism, considered that particular arrangements of lines and colours result in aesthetic experiences and consequently made such arrangements central to their art. The English art critic, Clive Bell, believed that “…lines and colours combined in a particular way [to produce] certain forms and relations of forms, stir our aesthetic emotions”; he did not specify what these particular combinations may be but argued that, “It need be agreed only that forms arranged and combined according to some *unknown and mysterious* laws do move us in a particular way and that it is the business of the artist to combine and arrange them that they shall move us” (Bell, 1914) (our emphasis). Thus, there is, in abstract paintings that arouse the aesthetic emotion, a mysterious and ineffable quality, just as there is in faces. Could that mysterious element also be represented in the form of decodable patterns in sensory areas of the visual brain that process lines and colours, just as happens in sensory face-processing areas when faces that arouse the aesthetic emotion are viewed? And would the emergence of decodable patterns in these sensory areas also correlate with the parallel emergence of decodable activity in medial frontal cortex, thus mirroring the pattern of brain activity during the experience of facial beauty? If so, an overall plausible interpretation would be that, whatever the mysterious qualities that endow stimuli, irrespective of their provenance, with the capacities of arousing the aesthetic emotion of beauty, they are represented in detectable patterns within the sensory areas that are specialized for processing the category of stimuli as well as decodable activity within the medial frontal cortex.

To avoid confusion, we begin by giving a brief terminological guide to the area of activation in medial prefrontal cortex. Although involving a common area, the exact location of activations there has varied across studies of the neural correlates of beauty, and different anatomical terms have been used to refer to the location in the literature, including medial orbitofrontal cortex (mOFC), medial prefrontal cortex (mPFC), ventromedial prefrontal cortex (vmPFC), and anterior cingulate cortex (aCC). Ishizu and Zeki (2011) addressed this point and suggested that the region of medial frontal activations related to aesthetic experiences be labelled ‘Field A1’, a functionally defined region which does not necessarily obey anatomical or cytoarchitectonic boundaries; field A1 has its centre at Montreal Neurological Institute (MNI) coordinates [−3 41 - 8] mm and a diameter of 15-17 mm. Therefore, in this paper we will refer to any activations within the medial frontal cortex which fall within mOFC, mPFC, vmPFC and aCC as Field A1. For this study, we expected that visual areas reported to have large concentrations of orientation-selective and colour selective cells will be active, namely, V1-V4 (including areas V3A and V3B) as well as areas in the intraparietal sulcus (Brouwer and Heeger, 2009, 2013; Cheadle and Zeki, 2014; Hubel and Wiesel, 1968; Montaser-Kouhsari et al., 2007; Zeki, S, Perry, RJ and Bartels, 2003; Zeki, 1978; Zeki and Stutters, 2013), *inter alia*. This leads us to conclude that the co-operative emergence of activity in both sensory areas and in field A1 constitutes the neural selection of objects with qualities that lead to the arousal of the aesthetic emotion and hence the experience of beauty. The experiments that we report here addressed questions based around these issues.

## METHODS

### Participants

18 healthy subjects (11 females, 7 males; ages 20-31 years, mean age 26.5 ± 3.2) participated in the study; all were right-handed and had normal or corrected-to-normal vision, all gave informed consent, and none was an artist or had art expertise. The experiment was approved by the ethical committee of Ludwig-Maximilians-Universität Munich (LMU) where the imaging experiments were conducted.

### Stimuli

The stimuli consisted of 120 images of abstract paintings. The images were obtained from one of our previous studies (Bignardi et al., 2020) and additional paintings were selected from stock image websites. To ensure that the set contained enough paintings belonging to the three categories of “not-beautiful”, “average”, and “beautiful”, a pilot study was undertaken in which 13 participants rated the paintings (Figure S3). The stimuli were scaled to 500×500 pixels, presented in their original colour, and were not cropped or modified. The task was programmed in the Presentation software package (Neurobehavioral Systems, Inc., Albany, CA). We excluded iconic paintings and schools, such as those of the De Stijl school, Russian Constructivists and Suprematists as well as works belonging to artists such as Barnet Newman and Rothko, which contain colours, oriented lines, and edges but which are easily recognizable as belonging to particular artists or schools. The set of stimuli used in this study is available as supplementary data.

### Procedure

Participants were presented with 120 images of abstract paintings inside the MRI scanner and asked to rate them on a scale from one to seven, one corresponding to not ‘beautiful at all’ and seven to ‘very beautiful’. Pressing the left button on a customized button box decreased the rating, while pressing the right button increased it, starting from the neutral rating of four at each trial. The task followed an event-related design in which the stimulus was presented for a duration of 2 s and participants had 4.5 s to respond (Figure 1). The scanning consisted of five functional magnetic resonance imaging (fMRI) runs, each containing 24 trials. This, in addition to structural scans, amounted to a total scan time of 45 minutes for each subject. After scanning, subjects were asked to complete a 15-minute questionnaire in which the same paintings were rated on familiarity, valence, arousal, and beauty.

**Figure 1.**
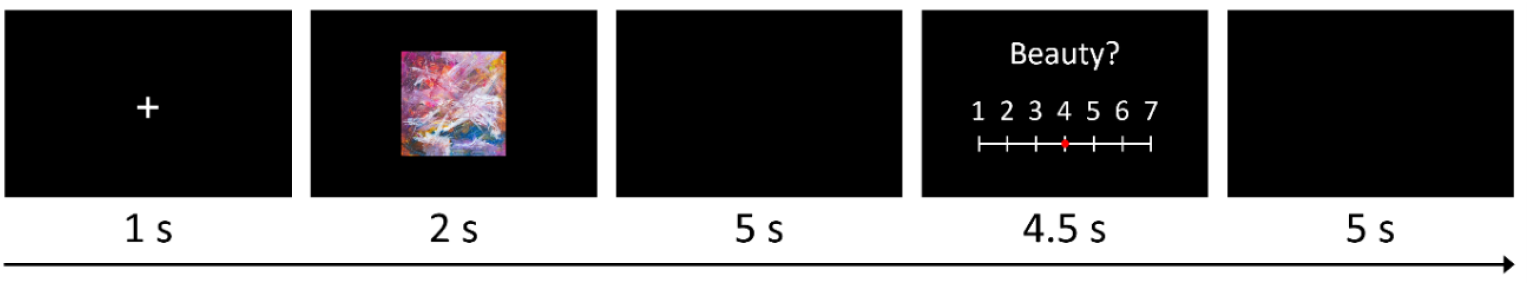
Experimental design. Each trial started with a 1 s fixation cross, followed by a 2 s stimulus presentation, which was followed by a 5 s blank screen. Subjects then rated the stimulus on aesthetic appeal within 4.5 s on a scale from 1 to 7 (1 – not beautiful at all, 7 – very beautiful). Responses were followed by a 5 s blank screen, amounting to a total trial time of 17.5 s. Each fMRI run consisted of 24 trials, yielding a total of 120 trials.

### Image acquisition

Brain images were collected at the University Hospital of the LMU on a 3.0 T Philips Ingenia scanner (Philips Healthcare, Best, The Netherlands). Structural images were acquired with a T1-weighted scan: repetition time (TR) = 9.74 ms; echo time (TE) = 5.66 ms; flip angle = 8°; matrix of 256×256; field of view = 256 mm; voxel size = 1×1×1 mm^3^. The blood oxygen level dependent (BOLD) signal was measured with a T2*-weighted Echo Planar Imaging (EPI) sequence: TR = 2500 ms; TE = 30 ms; flip angle = 90°; ascending acquisition; matrix of 80×80; voxel size = 3×3×3 mm^3^. A total of five fMRI runs were acquired. Field mapping data were also acquired using a dual-echo GRE sequence to assist with susceptibility distortion correction.

### Image preprocessing

The T1w image of each subject was skull-stripped using *optiBET* (Lutkenhoff et al., 2014), bias field corrected using the N4 tool (Tustison et al., 2010), and rigidly aligned, using flirt (Jenkinson et al., 2002), to the 1 mm MNI T1w brain template (Fonov et al., 2011) as a substitute for AC-PC alignment. This aligned image served as the anatomical reference for subsequent pre-processing of functional images. Additionally, the T1w image was normalised to the MNI template through affine and non-linear transformations (SyN algorithm) computed using ANTs (Avants et al., 2011). The rigidly aligned T1w image was passed to FreeSurfer (Fischl, 2012) to obtain a model of each subject’s cortical surface. Additional steps using tools from Connectome Workbench (https://www.humanconnectome.org/software) were applied to remap the surface of each subject to a common space, the ‘32k_FS_LR’ template. We specifically used these surfaces for each subject because they have the advantage of maintaining the native anatomy of the brain while offering a vertex-level matching between subjects. As a result, information from multiple subjects can be directly compared at each vertex. These surfaces were used for the multivariate analysis.

The first six volumes of the functional series were discarded to allow the scanner to reach steady state. This short period (15 s) was used to display instructions to participants reminding them of the task details. The remaining functional images were first corrected for motion and slice-timing differences using *SPM12* (http://www.fil.ion.ucl.ac.uk/spm/software/). The corrected images were then simultaneously corrected for geometric distortions (using the acquired field map) and aligned to the T1w image using FSL’s *epireg* tool (Greve and Fischl, 2009; Jenkinson et al., 2002), while maintaining the voxel size at 3×3×3 mm^3^. This produced the final fMRI time series images which were used in subsequent analyses.

### Image space and spatial smoothing

In this study, we analyse the data following the univariate and multivariate frameworks; both rely, in the first instance, on a subject-level (first level) model fitting using the general linear model (GLM). For the univariate analyses, the fMRI series of each subject were normalised to MNI space (at 3 mm) and spatially smoothed with a Gaussian kernel of a FWHM of 6 mm before running the first-level GLM.

For the multivariate analysis, the first-level GLM was performed on each subject’s data in native space without smoothing and the beta images (parameter estimates) were projected to the subject’s cortical surface (obtained from *FreeSurfer*). The surface-based beta maps were then used for the multivariate analysis. This was important to maintain the spatial specificity of the parameter estimates, which is crucial for the multivariate framework.

### Univariate analysis

Parametric analyses were performed to identify the brain regions in which activity increased (linearly or non-linearly) with beauty ratings. A standard GLM was fitted to the time series of each subject, with a single task effect (stimulus presentation), regressors that modelled the responses and rest periods, and six motion correction parameters as nuisance regressors. The beauty score given at each trial was used as an additional regressor to account for any variability in the BOLD signal that was not explained by the other regressors.

The parametric analyses were performed according to three models. The first was the classical linear parametric model which assumes a linear increase in BOLD signal with increasing beauty ratings. In this model, ‘not beautiful’ stimuli would be associated with the weakest activations and ‘very beautiful’ stimuli with the strongest. The second model assumed a V-shaped relationship between beauty ratings and brain activity, whereby ‘very beautiful’ and ‘not beautiful’ stimuli would lead to a similar level of activity, and ‘neutral’ stimuli would be associated with the weakest activity. The third model assumed a ‘checkmark-shaped’ relationship between beauty ratings and brain activity, whereby ‘very beautiful’ stimuli are associated with the highest activity, followed by ‘not beautiful’ stimuli, and finally by ‘neutral’ stimuli. Visual representations of these models are shown in Figure 3.

Categorical analyses comparing the ‘very beautiful’ condition (ratings of 6 and 7) to both ‘neutral’ (rating of 4) and ‘not-beautiful’ (ratings of 1 and 2) conditions were performed as well. To this end, we included all trials belonging to a given condition to conduct a robust GLM. These results are reported in the supplementary material.

After processing each subject’s time series at the first-level in *SPM12*, second-level modelling was performed through non-parametric permutation testing using the *SnPM* toolbox (Holmes et al., 1996; Winkler et al., 2014) as this type of group-level modelling for univariate analyses has been shown to be the most robust to false positives (Eklund et al., 2016). The first-level contrast images from all subjects were submitted to a one sample t-test and 5000 sign flipping permutations were performed to estimate the null distribution of the t-statistic at each voxel. The final statistical maps were created with a cluster-forming threshold of *p* < .001 and cluster-level FWE correction threshold of *p* < .05.

### Representational similarity analysis

To investigate whether the experience of abstract paintings as beautiful is associated with specific spatial patterns of neural activity, we used representational similarity analysis (RSA) (Kriegeskorte et al., 2008). We started by running a GLM for each subject in which each trial was treated as an independent condition, thereby generating a parameter estimate (beta) map for each trial. The beta maps corresponding to each subject’s 10 highest and 10 lowest rated paintings were selected (Figure S5) and projected to the brain surface. A whole-brain, surface-based searchlight analysis was performed using cortical patches with a 6 mm radius as regions of interest (ROIs). This allowed us to estimate the patterns of neural activity associated with these trials. For each searchlight ROI, the Pearson correlation distance, d, was calculated between the patterns associated with these trials as follows:

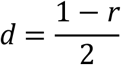

where *r* is the Pearson correlation coefficient, and the division by two was performed to rescale *d* to the range [0-1]. This was done to simplify the interpretation and visualisation of the metric: *d* = 0.0 corresponds to a full correlation between the neural patterns of two trials (i.e., *r* = 1.0); *d* = 0.5 corresponds to the absence of any correlation (i.e., *r* = 0.0); and *d* = 1.0 corresponds to the two trials having completely anti-correlated patterns (i.e., *r* = −1.0).

Once the Pearson distances were calculated for each pair of trials, neural representational dissimilarity matrices (RDMs) were generated for each subject to capture the (dis)similarity between pairs of trials in each searchlight ROI. The Pearson correlation distance is mainly sensitive to the spatial pattern of brain activity and is insensitive to the overall BOLD signal amplitude change in a given ROI (Kriegeskorte et al., 2008). Given that the aim here is to find a specific pattern of activity regardless of amplitude, the Pearson correlation distance is the preferred distance metric for our purposes (unlike, e.g., the Euclidean distance which would also record overall magnitude changes like the univariate framework).

To test whether the similarity was significant only for beautiful trials, the mean group RDM was calculated for each searchlight ROI, and these mean neural RDMs were then compared to a model RDM (Figure 4). The correlation between the neural and model RDMs was assessed with the Spearman rank correlation only using the elements in the lower triangle of the RDMs (excluding the diagonal), and statistical significance was determined by way of permutation testing. For each ROI of the searchlight, 5000 random permutations of the trial labels were generated to estimate the null distribution of the distance, *d*, and obtain a robust measure of statistical significance.

The first model RDM we tested (Figure 4) assumed a high similarity in the activity patterns associated with viewing ‘very beautiful’ paintings (i.e., *d* = 0.0), and no similarity when viewing paintings which were deemed ‘not beautiful’ or between the patterns of ‘very beautiful’ and ‘not beautiful’ paintings (i.e., *d* = 0.5). No pairs of trials were expected to have anti-correlated patterns (i.e., *d* = 1.0) as this would be a strong assumption to make. We also tested a second RDM model, which conversely assumed high similarity in activity patterns for ‘not beautiful’ paintings (Figure 4).

### Anatomical atlases

We referred to two atlases of cortical regions to label our results; the first was specifically used for the visual regions, and was based on a retinotopic mapping study (Wang et al., 2015); the second was the default one used by *FreeSurfer*, namely the Desikan-Killiany atlas (Desikan et al., 2006).

## RESULTS

### Behavioural results

The mean beauty rating for the abstract paintings was 3.97 (*sd* = 0.70). There was a low mean correlation of *r* = 0.16 (lowest Pearson *r* = −0.34, highest *r* = 0.61) when comparing each subject’s ratings to every other subject’s ratings, indicating that there was little agreement among subjects about each painting’s beauty score (Figure 2a). For comparison purposes, we reused behavioural data from a previous study on face beauty (Yang et al., 2022) and ran the same behavioural analysis. This revealed that the agreement among subjects is much higher for faces than it is for abstract art, with a mean correlation of *r* = 0.72 (lowest *r* = 0.42, highest *r* = 0.88), which is in accordance with previous studies (Bignardi et al., 2020).

**Figure 2.**
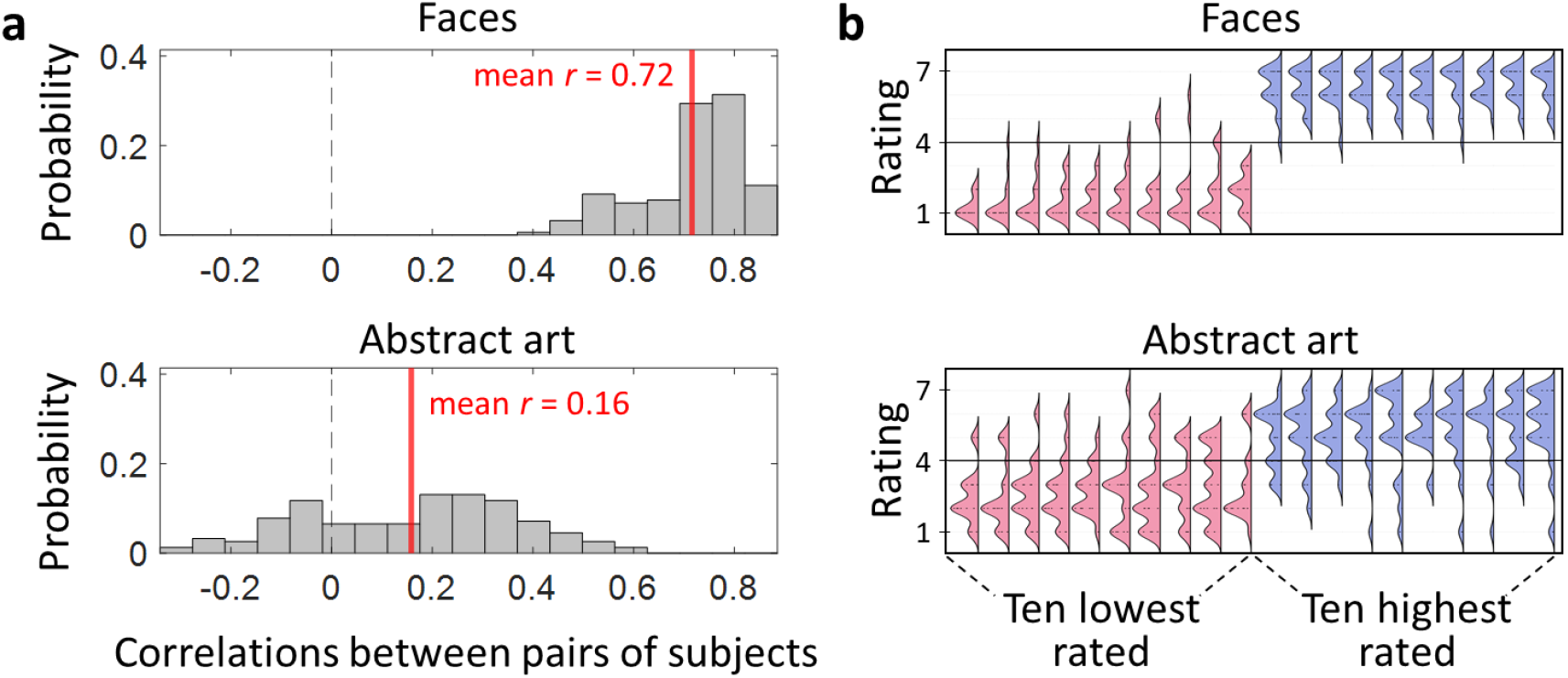
Behavioural results and comparison with data on face beauty. (a) Histograms showing the distribution of inter-subject correlations for beauty ratings given to a set of 120 faces (Yang et al., 2022) and 120 abstract art paintings (this study). For each study, we calculated the Pearson correlation coefficient between the beauty ratings of every pair of subjects, and here we plot the probability distributions of these correlations. The mean correlation score is indicated by the red lines. (b) Density plots showing the probability of ratings given to the 10 faces (Yang et al., 2022) and 10 abstract paintings (this study) with the best average (blue) and worst average (pink) scores. Each density plot was constructed using the scores given by all participants for each of these stimuli.

The great variability in the aesthetic ratings of abstract art is also well captured by the distributions of the ratings given by all subjects to the 10 abstract paintings and the 10 faces with the highest and lowest mean beauty scores (Figure 2b). With faces, the highest rated stimuli (on average) were predominantly given high ratings (higher than the neutral point of four), and those with the lowest mean beauty scores showed a similar trend with most subjects scoring them less than four. However, with abstract art, the highest rated stimuli still received low ratings (as low as one), and the lowest scoring paintings still received scores as high as seven. This indicates that there is greater universal agreement among subjects on the beauty of faces, and much less agreement on the aesthetic status of abstract art paintings. These findings sit well with the theory which proposes a distinction between biological and artifactual stimuli (Zeki and Chén, 2020).

### Univariate parametric activations

Several studies have reported that activity in field A1 increases linearly with the declared intensity of the experience of beauty, attraction, or desire, or that a more intense experience of beauty is associated with a categorically stronger activity in that region (Jacobsen et al., 2006; O’Doherty et al., 2003; Yang et al., 2022; Zeki et al., 2014). Yet some of the same reports, as well as others, have also pointed to a non-linear relationship between the level of the declared experience of beauty and BOLD signal changes in field A1, with neutral beauty ratings being associated with the weakest BOLD signal (Kawabata and Zeki, 2008; Martín-Loeches et al., 2014; Zeki et al., 2014). To further investigate this observation, we assessed the BOLD signal changes in our study according to three parametric models based on this previous literature.

#### Linear relationship with beauty

A linear parametric analysis of fMRI data with beauty as a modulator, that is, assuming a linear increase in brain activity as a function of increasing aesthetic appeal, revealed significant clusters in visual cortex (V1, V2, V3 and V4), but not field A1 (Table 1 and Figure 3).

**Table 1.** Results of the parametric fMRI analyses. Results of the three univariate parametric models assessing the relationship between BOLD activity and beauty ratings. The group-level analysis was carried out using permutation testing, with a cluster-forming threshold of *p* < .001 and FWE correction (*p*_clust-FWE_ < .05). These results and the models are visually represented in Figure 3.

| CLUSTER AND/OR REGION | VOXELS | P<sub>Clust-</sub><br>FWE | T | Coordinates (mm) |  |  |
| --- | --- | --- | --- | --- | --- | --- |
|  |  |  |  | x | y | z |
| *Linear parametric activations* |  |  |  |  |  |  |
| Visual cortex | 288 | 0.0196 |  |  |  |  |
| R V1 |  |  | 5.92 | 9 | -96 | 3 |
| L V2 and V3 |  |  | 4.05 | -12 | -90 | -18 |
| L V4 |  |  | 4.01 | -27 | -84 | -15 |
| *V-shaped parametric activations* |  |  |  |  |  |  |
| Superior frontal gyrus | 334 | 0.0156 | 7.31 | 3 | 51 | 36 |
| L lateral orbitofrontal cortex | 239 | 0.0230 | 5.62 | -39 | 27 | -15 |
| R lateral orbitofrontal cortex | 171 | 0.0336 | 4.87 | 27 | 18 | -18 |
| *Checkmark-shaped parametric activations* |  |  |  |  |  |  |
| Prefrontal cortex | 1644 | 0.0020 |  |  |  |  |
| L posterior OFC and ventral striatum |  |  | 6.22 | -18 | 9 | -24 |
| Field A1 |  |  | 6.09 | -3 | 48 | 0 |
| R posterior OFC and ventral striatum |  |  | 5.89 | 21 | 15 | -21 |
| Visual cortex | 96 | 0.0496 |  |  |  |  |
| R V1 |  |  | 4.47 | 15 | -87 | 3 |
| R V1 |  |  | 4.16 | 9 | -84 | -3 |
| R V2 and V3 |  |  | 4.04 | 21 | -81 | -3 |

**Figure 3.**
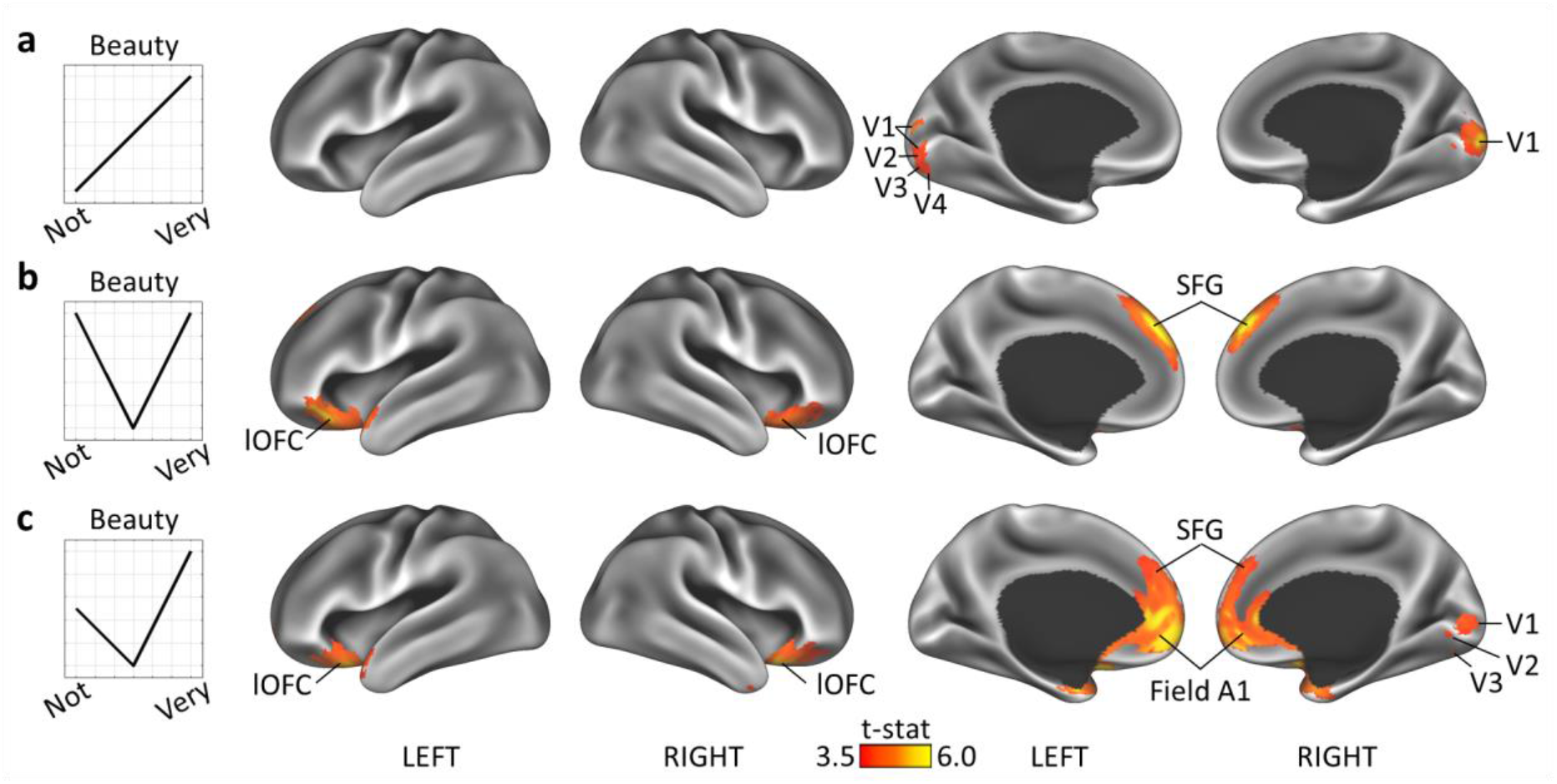
Parametric BOLD activity during aesthetic experiences. BOLD activity during a beauty rating task was assessed according to three parametric models. (a) Model 1 assumed a linear increase in activity with increasing beauty ratings. The only cluster revealed by this model was in visual cortex (V1, V2, V3 and V4). (b) Model 2 assumed a V-shaped relationship between beauty ratings and brain activity, whereby ‘very beautiful’ and ‘not beautiful’ stimuli would lead to a similar level of activity, and ‘neutral’ stimuli would be associated with the lowest activity. This model revealed clusters in lateral orbitofrontal cortex (lOFC) and the superior frontal gyrus (SFG), bilaterally. (c) Model 3 assumed a ‘checkmark-shaped’ relationship between beauty ratings and brain activity, whereby ‘very beautiful’ stimuli are associated with the highest activity, followed by ‘not beautiful’ stimuli, and finally by ‘neutral’ stimuli. This model was associated with the largest cluster of activity in field A1 of medial prefrontal cortex (aCC and mOFC), and in the ventral striatum (not shown here), as well as the SFG and lOFC. All results are based on non-parametric permutation testing (pclust-FWE < .05). MNI coordinates are shown in Table 1.

#### Deviation from neutrality: V-shaped model

A V-shaped parametric model, assuming an equal increase in brain activity in either direction (‘not beautiful’ or ‘very beautiful’) compared to neutrality, revealed that a large portion of the superior frontal gyrus (SFG) and bilateral lateral orbitofrontal cortex (lOFC) showed increased activity with more extreme beauty judgments, both toward the high and low ends of the scale (Table 1 and Figure 3).

#### Deviation from neutrality: Checkmark-shaped model

Finally, the checkmark-shaped parametric model also assumed an increase in brain activity in either direction (‘not beautiful’ or ‘very beautiful’) compared to neutrality, but to a different extent: activity related to ‘not beautiful’ stimuli was expected to be weaker than that related to ‘very beautiful’ stimuli. This model revealed strong activations in field A1, bilateral lOFC, SFG and visual cortex (V1, V2 and V3) (Table 1 and Figure 3).

### Representational similarity analysis

A whole-brain searchlight representation similarity analysis (RSA) using the Pearson correlation distance revealed clusters in visual areas with common patterns in response to beautiful stimuli. More specifically within this RSA context, we compared the representational dissimilarity matrices (RDMs), which contain the dissimilarity scores between pairs of trials, to a model RDM that assumed similar patterns only for ‘very beautiful’ stimuli. Spearman correlations between the neural and the model RDM revealed the following visual regions with significant correlations after permutation testing, according to the Wang et al. (2015) atlas of visual regions: left V1, right V2v/V3v, bilateral V3, left VO2 (anterior to V4α), left V7/IPS0, bilateral anterior fusiform gyrus and left SFG (Figure 4 and Table S2). Model 2, which assumed similar patterns only for the ‘not beautiful’ stimuli, only correlated significantly with the neural RDM of anterior right V1.

**Figure 4.**
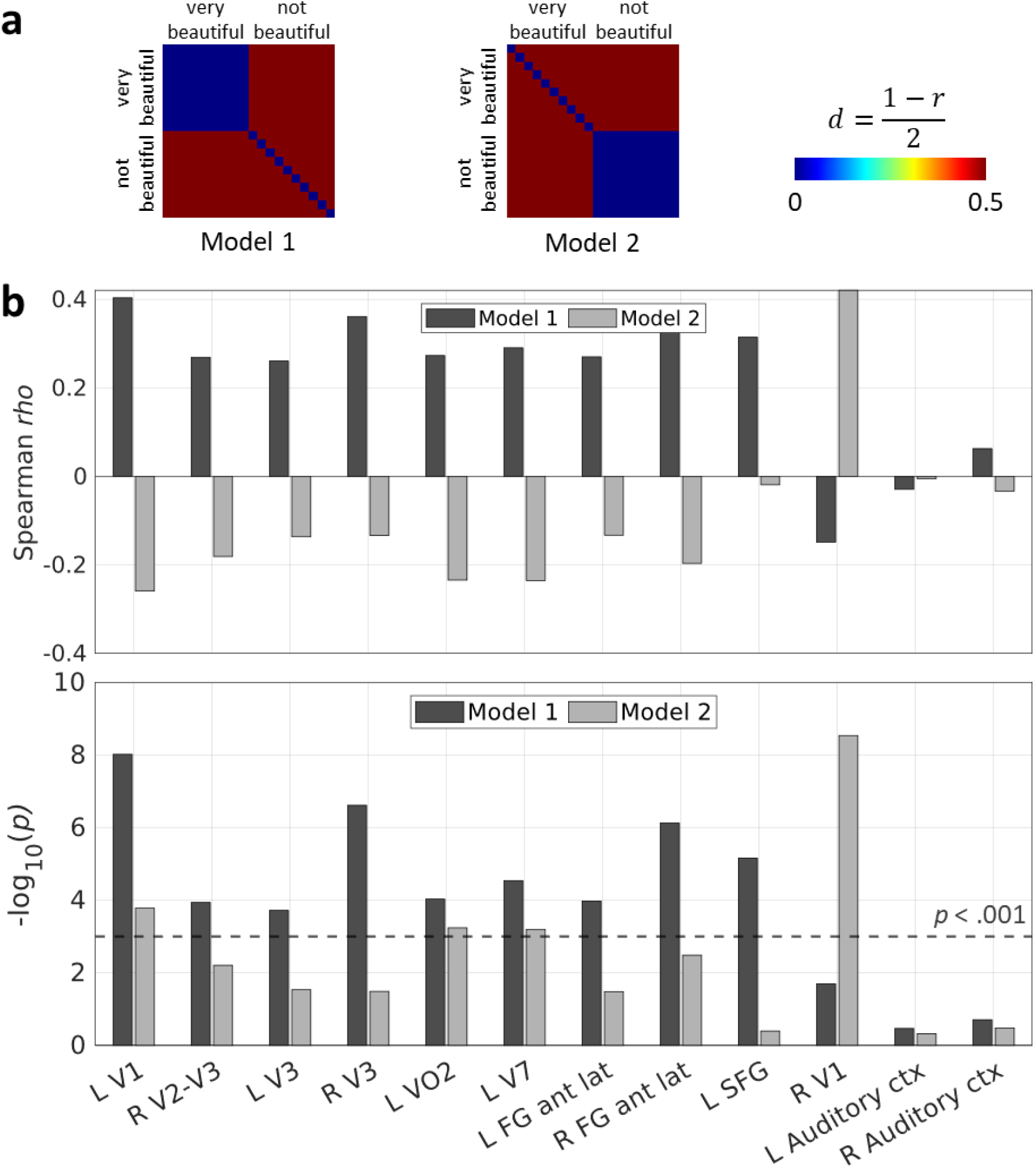
Results of the representational similarity analysis. a) The model representational dissimilarity matrices (RDMs) used for RSA. The RDMs represent the relationship between pairs of trials based on the Pearson distance metric, calculated based on the displayed equation. Smaller distances equal greater similarity between pairs of trials. Model 1 assumes that only ‘very beautiful’ stimuli share similar neural patterns, while Model 2 assumes that only ‘not beautiful’ stimuli share similar patterns. b) The top bar plot shows the Spearman rank correlation between the neural RDMs for a set of regions and the two model RDMs obtained through a searchlight analysis. The bottom bar plot displays the significance of each reported correlation. Most of the regions that were positively and significantly correlated with Model 1 were in visual cortex. The dashed line represents the threshold for significance at p < .001. For comparison, we also included a region which is not expected to be involved in the task, namely the primary auditory cortex, which indeed did not show any correlation with either model. MNI coordinates are shown in Table S2.

The results obtained from the two models indicate that the patterns in visual cortex are indeed specific to the ‘very beautiful’ category. Figure 4 shows that most visual regions correlated positively with Model 1 and negatively with Model 2. This is expected to some degree given that the two models assume almost opposite trends but the two need not be fully anti-correlated. Figure 4 also shows that a control region which is not expected to be involved in the task, namely primary auditory cortex, was not correlated to either model.

## DISCUSSION

### Biological vs. artifactual beauty

We enquired into whether the experience of beauty derived from viewing abstract works of art – consisting of arbitrary assemblies of lines and colours – leads to broadly similar neural activity as that resulting from viewing beautiful human faces based on previous studies. Though both can arouse an experience of beauty, the two categories of stimuli differ significantly. While the experience of facial beauty is mediated through inherited or rapidly acquired concepts and is resistant to revision through peer opinion (Chen and Zeki, 2011; Glennon and Zeki, 2021), the experience of works created through human agency, in which we include abstract works of art, is much less resistant to revision in light of peer opinion and is probably not interfaced through any known inherited brain concept (Bignardi et al., 2020; Zeki, 2009; Zeki and Chén, 2020). Yet the fact that they can both arouse the “aesthetic emotion”, or the experience of beauty, implies that aesthetic experiences aroused through a combination of lines and colours in abstract paintings may engage broadly similar brain mechanisms. It thus became interesting to learn whether the experience of beauty in abstract works of art would result in decodable activity within sensory visual areas on the one hand and within field A1 of medial prefrontal cortex on the other.

When we place abstract art in the artifactual category, we do not mean to imply that the constituents that go into making up abstract art may not be the result of inherited physiological mechanisms. It is almost certain, for example, that orientation selectivity is inherited (Hubel and Wiesel, 1977); but the assembly of lines and colours in different works is not. This is distinct from faces where the constituent elements usually have to take their correct place in an overall composition in order to be recognised as a face. Consistent with previous results (Bignardi et al., 2020), our present behavioural evidence shows that there is much greater variability in assigning particular abstract works to particular aesthetic categories, making of the experience of beauty in abstract art a more subjective one than the experience of facial beauty.

Despite these differences, a similar strategy is used by the brain for the two categories; both involve the emergence of decodable patterns in sensory areas; with facial beauty the decodable activity was in the sensory areas known to be critical for the perception of faces (Yang et al., 2022), while for beautiful abstract art it was in sensory visual areas known to contain large concentrations of orientation selective and chromatic cells. In addition, there is increased activity in field A1 as revealed by the univariate analyses. However, there remains one important difference: whereas Yang et al. (2022) found distinct patterns in field A1 for beautiful faces, we did not find such patterns in field A1 for beautiful abstract art. Therefore, these results suggest that a principal difference between facial and abstract beauty is that for faces (biological beauty), a pattern emerges in both the sensory areas and field A1, but for abstract art (artifactual beauty), a pattern emerges in the sensory areas only. We present this explanation tentatively and with diffidence because it seems overwhelmingly simple, and we await future studies to confirm it. Our hesitation can also be traced to the fact that the study of Vessel et al. (2019) reported decodable activity in field A1 in response to both beautiful buildings (artifactual) as well as to beautiful faces (biological). However, our method (RSA with the Pearson correlation distance) is more suitable for handling the specific question addressed in this study, that of detecting patterns of neural activity, as opposed to using multivariate pattern analysis (MVPA) which is sensitive to both patterns or overall amplitude of activity (Kriegeskorte et al., 2008).

### The selective function of sensory areas

The sensory areas of the visual brain, in addition to the sensory functions attributed to them – of processing visual stimuli according to their specialisations – also have the function of selecting stimuli according to certain arrangements of their constituent elements, endowing them with aesthetic appeal. Clive Bell (1914) referred to such arrangements, dictated by “unknown laws” as “Significant Forms”; we prefer to generalise the term and refer instead to “Significant Configurations” (Zeki, 2013); the former term sits uneasily with, for example, harmonious colours or beautiful facial expressions, while the latter term does not.

Our results, and those of Yang et al. (2022), show that the so-called visual “sensory” areas cannot be mere passive recipients of signals related to their specialities. Rather, the ineluctable conclusion seems to be that these sensory areas are also able to classify stimuli according to their aesthetic appeal. This seems to be true regardless of whether the viewed stimuli belong to the biological category (faces) or the artifactual one (abstract art). This is because decodable patterns emerge in the sensory areas only with stimuli that are experienced as beautiful, regardless of category, and it is only when such patterns emerge that there is, as a correlate, activity in field A1.

It is tempting to suggest that it is the emergence of such decodable patterns in sensory areas that engages field A1. We do not do so and deliberately use the term ‘as a correlate’ instead because, in the absence of temporal studies, we cannot exclude the possibility that field A1 is activated prior to the sensory areas and that the latter are only activated through feedback. A precedent for this may be found in studies which have shown, for example, that activity in the amygdala precedes activity in face-processing sensory areas when fearful stimuli are viewed (Méndez-Bértolo et al., 2016). Whatever the temporal relationship in activity between the sensory areas and field A1, we cannot escape the conclusion that, in addition to registering the characteristics of the stimulus, the sensory areas are also involved – either before, simultaneously with, or after activation of field A1 – in ordering the stimuli according to criteria that give them an aesthetic status. They are therefore involved in a selective process.

One criticism may be that certain low-level features, such as colour or brightness, may be more abundant in beautiful paintings than in not beautiful ones, thereby driving the increased response in visual cortex, instead of beauty itself (Iigaya et al., 2021). However, we believe that this point is circumvented by the variance in aesthetic ratings that we observed in our cohort (Figure 2): what was regarded as beautiful by some was regarded as not beautiful by others. Because of this, there was no fixed beauty condition that applied across subjects, which means that when making any comparison, a given painting is not consistently present in any category. Further, previous neuroimaging studies have specifically assessed the neural correlates of aesthetic judgement and judgement of other features of visual stimuli such as brightness and symmetry, and found that the network of orbitofrontal (medial and lateral) regions that we report is more involved in aesthetic judgements (Ishizu and Zeki, 2013; Jacobsen et al., 2006).

### Co-activity between sensory areas and field A1 is the basis of the experience of beauty

Our second conclusion is that it is only under conditions when decodable patterns of activity emerge in sensory areas, with activity in field A1 as a correlate, that the observer experiences stimuli as beautiful. We compared the neural activity associated with the perception of works of abstract art at three levels of aesthetic experience – ‘very beautiful’, ‘neutral’, and ‘not beautiful’. Our results showed that visual cortex is more engaged when the declared experience of beauty increases, and this engagement is further supported by the emergence of specific neural patterns in visual cortex. Moreover, we observed important differences within the medial prefrontal region: beautiful stimuli fully engaged field A1, while the other categories did not, or did so to a limited degree. Indeed, direct comparisons between the ‘very beautiful’ category and the other categories (Figure S7 and Figure S8), as well as the parametric analysis of brain activity according to the ‘checkmark-shaped’ model (Figure 3), showed strong activations in a large swath of medial frontal cortex in a region that included field A1, among others. This indicates that field A1 in the medial prefrontal region is involved in assigning a positive aesthetic attribute to a stimulus, or in processing the reward related to that stimulus, though we do not intend to imply that that is its only function. Therefore, it is the unique combination of increased activity in sensory cortex according to certain configurations and increased activity in field A1 that underlies the experience of beauty.

The activity in field A1 itself also raises interesting questions. Whether the same or different sub-regions of it are active with different aesthetic experiences is not clear (see also Pegors et al. (2015)). Hence, one pointer to future work that emerges from these studies is the importance of detailing the pattern of anatomical connectivity within field A1 and between it and other areas in both the sensory and frontal cortices. In light of our present studies and those of others, we also suggest that the boundaries of A1, as defined by Ishizu and Zeki (2011), be expanded, especially ventrally, to include other areas implicated in the experience of beauty reported in other studies (e.g., Tsukiura and Cabeza, 2011). We propose that field A1 should have a diameter of around 30 mm with the same central coordinates. We believe that the variability in reported activations is largely caused by (poor) fMRI signal quality in that region of the brain, especially across studies using different scanners and acquisition parameters (Weiskopf et al., 2006).

Activity in other prefrontal areas, such as the lateral orbitofrontal cortex (lOFC) and the superior frontal gyrus (SFG), also showed up consistently when subjects gave non-neutral ratings to the abstract art stimuli, indicating that these regions are also involved in aesthetic judgement. However, the exact role of these regions in the context of neuroaesthetics is yet to be confirmed. For example, are these regions revealed by our analyses because they play a general role in judgement, or could their role be more specific within beauty tasks? Previous research suggests that general involvement in judgement may be the driving factor (Elliott et al., 2000; Ishizu and Zeki, 2013; Kringelbach, 2005).

### The demonstration of experience-dependent connections in the human brain

Over a quarter of a century ago, Crick and Jones (1993) considered it “intolerable” and “shameful” that we do not have as much information about human neuroanatomy as we do about that of monkey; they were in a search for anatomical techniques that can be used in the post-mortem human brain, based around tract-tracing; this, they hoped, might reveal a connectivity pattern in the human brain which could come close to the anatomical tracing methods used in live monkeys. Since then, many advances have been made in the study of the human brain’s anatomy and function owing to advances in non-invasive brain imaging techniques, and these are of two kinds.

The first revolves around diffusion MRI and tractography which, together, have advanced our knowledge of the brain’s anatomical connectivity. They have demonstrated, for example, the existence of direct anatomical connections between the posterior occipital cortex and the prefrontal cortex in the human brain, and have linked them to conscious visual processing and face perception (ffytche and Catani, 2005; Forkel et al., 2014; Rokem et al., 2017). These studies (and others) have established that the polar occipital cortex is directly connected with the lateral and polar frontal cortex, but no direct connections have been established with the medial anatomical areas overlapping field A1. Despite these advances, the descriptions of the exact terminations of these connections are vague with respect to the many specific visual areas of the occipital lobe. It is hard to learn from what is available which visual areas such as V3, V3A, V4, FFA, or OFA are connected with which part of the frontal cortex.

The second approach is illustrated by the present results and those of Yang et al. (2022), among many others, which demonstrate that connections, whether direct or indirect, between the areas enumerated above do exist in the human brain but are currently only demonstrable during certain experiential states – in our instance during the experience of beauty. We refer to these as *experience-dependent connections in the human brain* which, otherwise, remain occult. The results also show that our experience dependent connections are selective, in that parallel activity in sensory areas and in field A1 depends upon the nature of the stimulus that elicits the experience of beauty.

We use the examples provided in this study and in that of Yang et al. (2022) although of course any number of previous fMRI studies could serve our purpose just as well. We also do not mean that these connections can only become evident with the experience of beauty; other affective experiences may also render them visible. For example, in a previous study (Ishizu and Zeki, 2014) we showed that ambiguous visual stimuli, in addition to engaging visual areas such as V1 and V3, also engage the mid-cingulate cortex, which had been implicated in conflict monitoring (Botvinick et al., 2001), though no direct connections between these visual areas and the mid-cingulate cortex have been anatomically demonstrated. Such connections also exist in other domains, such as that of motor learning (Laureys et al., 2001). But it is important to emphasize that there are such highly specific state-dependent connections that become demonstrable with specific aesthetic experiences.

### Limitations

There may be certain limitations to our study. For instance, we used RSA as a method for multivariate analysis, and this method relies on the distribution of activity across voxels or vertices. This, of course, opens the results to variability with a different imaging resolution. Perhaps higher resolution in future imaging experiments conducted at higher field strength could reveal patterns in other areas and help refine the ones we found in our study. Further, as with many other studies similar to ours, we based our results on a relatively small sample (18 subjects), and therefore future studies with larger samples would be valuable to see whether our findings can be replicated. The effects we demonstrate would serve as a valuable basis for others who wish to perform a formal power analysis prior to running such replication studies. Lastly, in this study our main focus was the experience of beauty. Given that the 7-point rating scale went from ‘not-beautiful’ to ‘very-beautiful’, one may argue that the ‘not-beautiful’ and ‘neutral’ categories are fairly similar in evoked experience. Therefore, we cannot make any claims about negative aesthetic judgements, such as ugliness. We hope to address this in future studies.

### Conclusion

We have demonstrated that the experience of beauty derived from abstract art leads to specific patterns of neural activity in a set of visual regions involved in processing orientations and colours, as well as to activity in medial frontal regions involved in affective processing and decision-making. We have further demonstrated that the connection between these geographically distant regions is specific, in that it is only established during the experience of works of art which the subjects deemed beautiful (in our experimental context). Our results, taken in conjunction with those of Yang et al. (2022), bring us closer to understanding the neural determinants of the experience of beauty; they also bring us a little, but not much, closer to understanding the nature of beauty itself. Finally, and with timidity, we suggest that a difference between the neural determinants of facial (biological) and abstract (artifactual) beauty may lie in the emergence of neural patterns in field A1.

## ACKNOWLEDGEMENTS

This work was supported by the Leverhulme Trust grant RPG-2017-341 to Semir Zeki. We thank Professor Ernst Pöppel for facilitating these experiments and Dr. Taoxi Yang for expert advice on several aspects of the study.

## AUTHOR CONTRIBUTIONS

S. Zeki conceived of the experiment; A. Beyh and S. Rasche conducted the experiments, analysed the data and, together with S. Zeki, wrote the manuscript; M. Paolini made the facilities at Munich available and made critical comments on the manuscript.

## DATA AVAILABILITY STATEMENT

The data of this study is available on request from the corresponding authors.

## LIST OF ABBREVIATIONS

aCC: anterior cingulate cortex
AC-PC: anterior commissure-posterior commissure
BOLD: blood oxygen level dependent
EPI: Echo Planar Imaging
FFA: fusiform face area
fMRI: functional magnetic resonance imaging
GLM: general linear model
GRE: gradient echo
lOFC: lateral orbitofrontal cortex
MNI: Montreal Neurological Institute
mOFC: medial orbitofrontal cortex
mPFC: medial prefrontal cortex
MVPA: multivariate pattern analysis
RDM: representational dissimilarity matrix
ROI: region of interest
RSA: representational similarity analysis
SFG: superior frontal gyrus
TR: repetition time
TE: echo time
vmPFC: ventromedial prefrontal cortex

