## Supplementary material for "The neural determinants of abstract beauty"

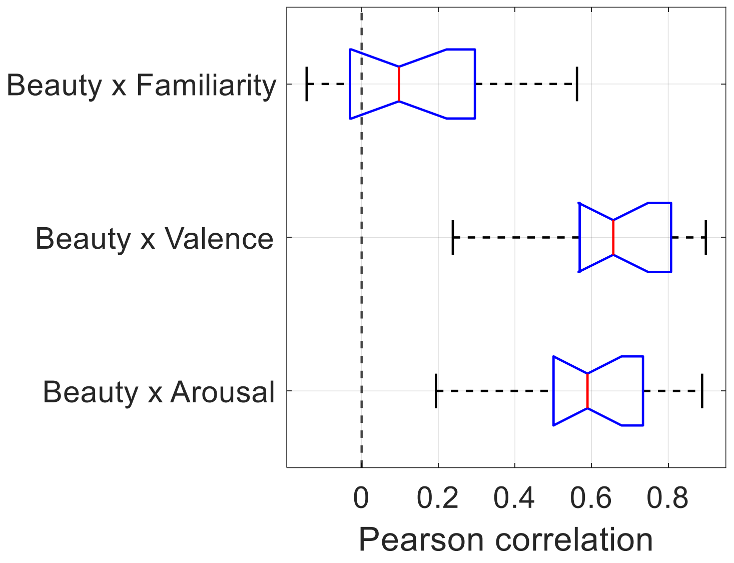


**Figure S1. Correlations between beauty and arousal, valence, and familiarity.**

For each subject (N = 18), the beauty scores of the 120 abstract art stimuli used in this study were correlated with their familiarity, valence, and arousal scores obtained in a post-scanning questionnaire. These box plots represent the distribution of the resulting Pearson correlation scores.

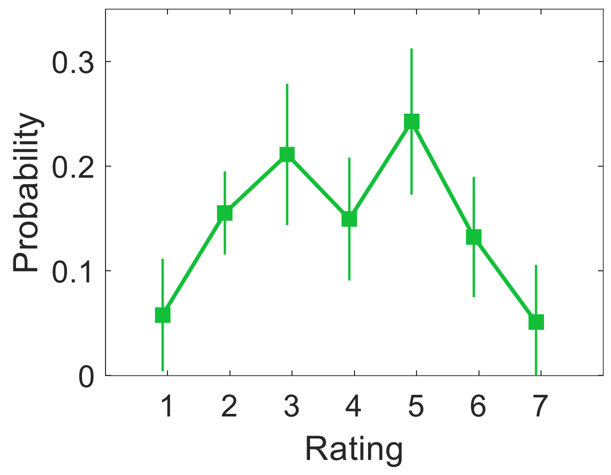


**Figure S2. Distribution of beauty ratings during the fMRI task.**

Probability distribution of beauty ratings given for the 120 abstract art stimuli used in this study. These ratings were given by subjects during the fMRI paradigm.

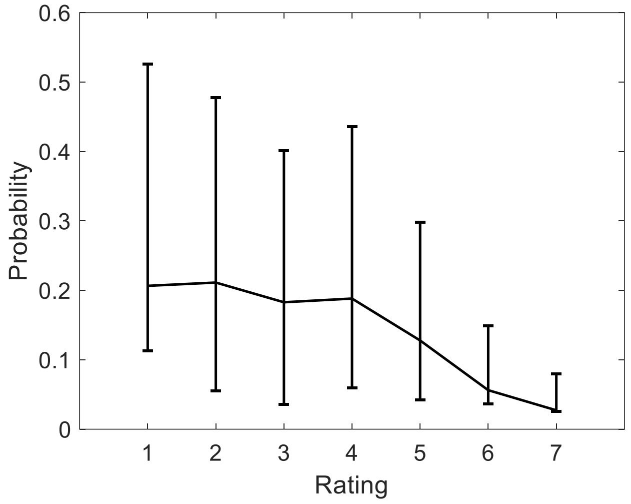


**Figure S3. Distribution of beauty ratings in the pilot study.**

Probability distribution of beauty ratings given in the pilot study for 120 abstract art paintings. This was done psychophysically on 13 subjects. The low number of high ratings led us to include more beautiful paintings (rated by the same subjects) in the final set of stimuli to ensure a more balanced distribution for the actual task.

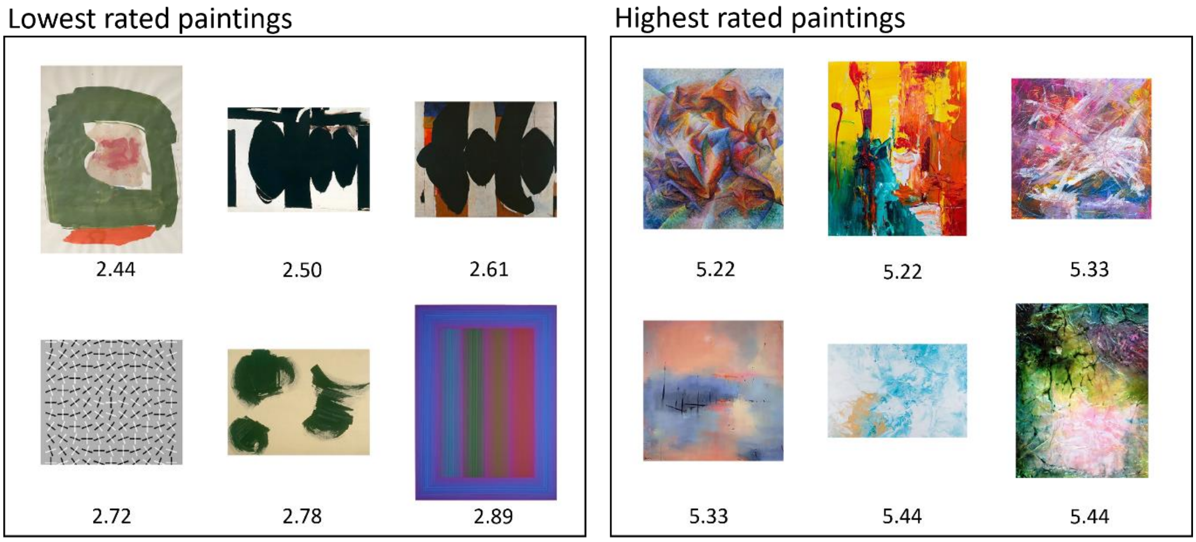


**Figure S4. Lowest and highest rated paintings on average.**

The six highest and six lowest rated paintings, on average, are shown here. As this is the average score, these paintings were not necessarily included in activity comparisons made between the 'not beautiful' and 'beautiful' categories, since the comparisons relied on each subject's own opinion. This is especially evident in Figure 2 in the manuscript which clearly shows a very large disagreement among subjects about the beauty of these paintings.

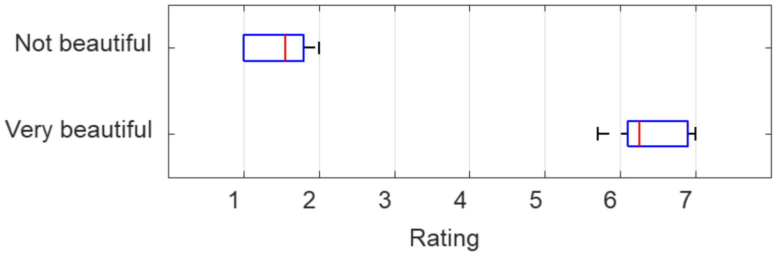


**Figure S5. Beauty scores of the beauty categories defined for RSA.**

The box plots in this figure represent the distribution of the beauty scores based on which two beauty categories were defined, with 10 trials in each, for representational similarity analysis (RSA): ‘not beautiful’ and ‘very beautiful’. These categories were defined for each subject individually based on their own scores, and this is what the RSA was based on. The mean score of each subject (N=18) was computed per category and the distribution of these mean scores is shown here. The median score of the ‘very beautiful’ category was 6.3 and that of the ‘not beautiful’ category was 1.6.

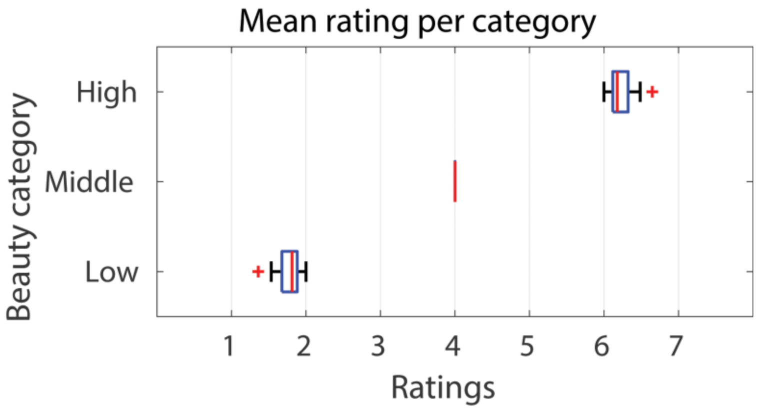


**Figure S6. Beauty scores of the beauty categories defined for the categorical analyses.**

The box plots in this figure represent the distribution of the beauty scores based on which three beauty categories were defined for the supplementary categorical univariate analyses (Figure S7 and Figure S8): ‘not beautiful’, ‘neutral’, and ‘very beautiful’. These categories were defined for each subject individually based on their own scores. The mean score of each subject (N=18) was computed per category and the distribution of these mean scores is shown here. The median score of the ‘very beautiful’ category was 6.2, that of the ‘neutral’ category was 4.0, and that of the ‘not beautiful’ category was 1.8.

| CLUSTER AND/OR REGION | VOXELS | $P_{Clust-FWE}$ | T | Coordinates (mm) x | y | z |
|---|---|---|---|---|---|---|
| ***Very beautiful > Not beautiful*** | | | | | | |
| R V1 | 109 | 0.0488 | 5.10 | 6 | -96 | 0 |
| Medial prefrontal lobe and caudate | 136 | 0.0422 | | | | |
| Anterior cingulate cortex (aCC) | | | 4.49 | 0 | 30 | 0 |
| R Caudate | | | 4.49 | 9 | 21 | 3 |
| aCC | | | 4.14 | 6 | 30 | 9 |
| ***Very beautiful > Neutral*** | | | | | | |
| Prefrontal cortex and striatum | 2083 | 0.0004 | | | | |
| Lateral orbitofrontal cortex (lOFC) | | | 6.89 | -33 | 18 | -21 |
| Field A1 (aCC) | | | 6.80 | -3 | 48 | -3 |
| Field A1 (right posterior OFC) | | | 5.70 | 18 | 12 | -18 |
| Visual cortex | 299 | 0.0154 | | | | |
| R V1 | | | 5.11 | 18 | -87 | 3 |
| R V3v | | | 4.68 | 21 | -81 | -3 |
| R Lateral occipital cortex (LO1) | | | 4.58 | 27 | -84 | 6 |

**Table S1. Results of the categorical fMRI analysis.**

Categorical univariate analyses revealed significantly higher activity in the visual cortex, field A1, lateral orbitofrontal cortex (lOFC), and the striatum during the experience of ‘very beautiful’ compared with ‘not beautiful’ or ‘neutral’ abstract art. Note that the size of the prefrontal cluster in the ‘very beautiful > neutral’ comparison is very large (2083 voxels). This cluster covers a large swath of bilateral lOFC and mOFC, and the bilateral striatum including the nucleus accumbens, which may not be clear by simply examining the peak coordinates. These results are shown on the cortical surface in Figure S7 and Figure S8.

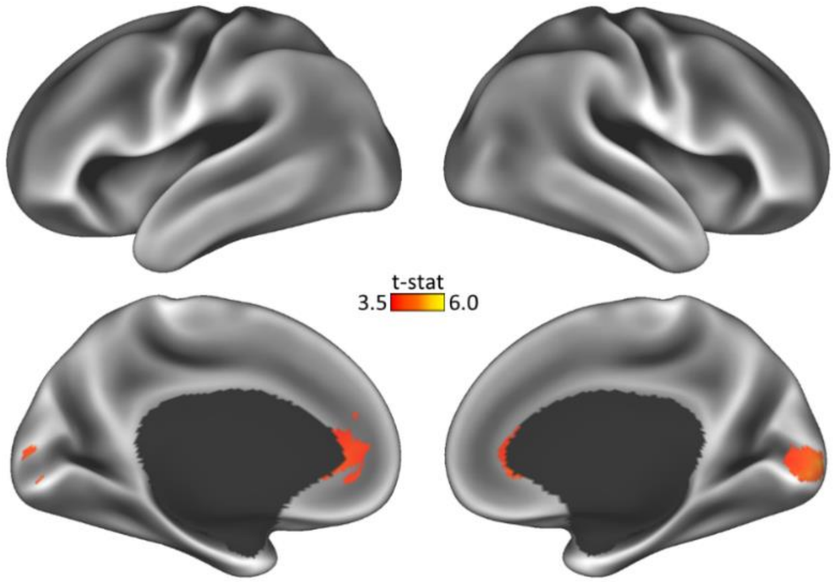


**Figure S7. Categorical analysis of ‘very beautiful’ > ‘not beautiful’ abstract art.**

Visual summary of the cortical locations where ‘very beautiful’ (scores of 6 and 7) abstract art stimuli elicited stronger activations compared to ‘not beautiful’ (scores of 1 and 2) stimuli. The main activations were in primary visual cortex (V1) and in field A1 (specifically, anterior cingulate cortex). Additional subcortical activations include the head of the caudate (not shown here).

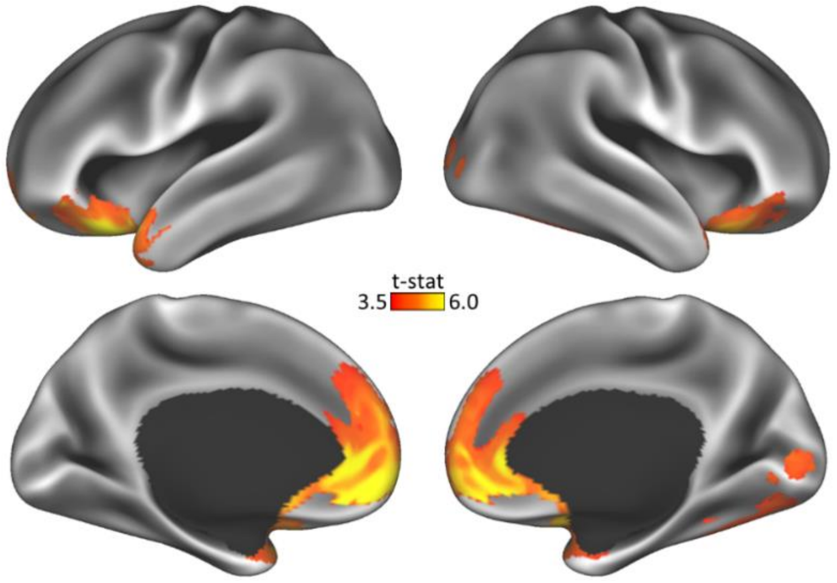


**Figure S8. Categorical analysis of ‘very beautiful’ > ‘neutral’ abstract art.**

Visual summary of the cortical locations where ‘very beautiful’ (scores of 6 and 7) abstract art stimuli elicited stronger activations compared to ‘neutral’ (score of 4) stimuli. The main activations were in visual cortex, field A1, lateral orbitofrontal cortex (lOFC), the superior frontal gyrus, and the striatum.

| REGION | *Model 1* $r_S$ | *Model 1* $-log_{10}(p)$ | *Model 2* $r_S$ | *Model 2* $-log_{10}(p)$ | MNI Coordinates (mm) | | |
|---|---|---|---|---|---|---|---|
| | | | | | x | y | z |
| L V1 | 0.4 | 8 | -0.26 | 3 | -7 | -97 | -5 |
| R V2-V3 | 0.27 | 3 | -0.18 | 2 | 19 | -76 | -15 |
| L V3 | 0.26 | 3 | -0.14 | 1 | -17 | -97 | 23 |
| R V3 | 0.36 | 6 | -0.13 | 1 | 29 | -90 | -6 |
| L VO2 | 0.27 | 4 | -0.23 | 3 | -30 | -61 | -9 |
| L V7 | 0.29 | 4 | -0.24 | 3 | -31 | -76 | 21 |
| L FG ant lat | 0.27 | 3 | -0.13 | 1 | -41 | -50 | -22 |
| R FG ant lat | 0.35 | 6 | -0.2 | 2 | 48 | -43 | -27 |
| L SFG | 0.31 | 5 | -0.02 | 0 | -21 | 35 | 50 |
| R V1 | -0.15 | 1 | 0.42 | 8 | 15 | -77 | 4 |
| L Auditory ctx | -0.03 | 0 | -0.01 | 0 | -42 | -28 | 11 |
| R Auditory ctx | 0.06 | 0 | -0.03 | 0 | 42 | -23 | 12 |

**Table S2. Results of the multivariate fMRI analysis with RSA.**

Representational similarity analysis (RSA) revealed significant pattern similarity in various areas in the visual cortex when 'very beautiful' stimuli were grouped together. We also found one location, namely right V1, in which significant pattern similarity appeared when 'not beautiful' stimuli were grouped together. The data in this table are represented in the main text in Figure 4.
